# Natural genetic variation reveals divergent transcriptomic responses to hyperoxia in two *Chlamydomonas reinhardtii* ecotypes

**DOI:** 10.64898/2026.07.09.737578

**Authors:** Joshua A. Temple, Peter G. Neofotis, Ben F. Lucker, Jacob D. Bibik, David M. Kramer, Daniela Strenkert

**Affiliations:** Department of Plant Biology, Michigan State University, East Lansing, MI, United States; MSU- DOE Plant Research Laboratory, Michigan State University, East Lansing, MI, United States; Department of Biology, Virginia Military Institute, Lexington, Virginia 24450; MelaTech, LLC, 1 N Haven St, Baltimore, MD 21224; Department of Biochemistry, Michigan State University, East Lansing, MI, United States; Jan Ingenhousz Institute, Wageningen, The Netherlands

**Keywords:** photosynthesis, hyperoxia, transcriptome, oxidative stress, reactive oxygen species, green algae

## Abstract

Green algae must continuously balance resource availability to maintain photosynthetic performance. The O_2_:CO_2_ ratio is a key determinant of their metabolic mode. Under hyperoxia or low CO_2_, many algae induce a carbon concentrating mechanism (CCM). In the model green alga *Chlamydomonas reinhardtii*, the CCM relies on a pyrenoid, a specialized microcompartment that elevates CO_2_ around rubisco. While ambient CO_2_ acclimation is well-studied, responses to hyperoxia remain poorly understood, despite its frequent occurrence in nature under high light. Using controlled bioreactors, we exposed two diverse *Chlamydomonas* ecotypes, CC1009 and CC2343, to 95% oxygen to analyze time-dependent, genome-wide transcriptomic and phenotypic changes. Both ecotypes induced CCM genes, but they exhibited distinct molecular and physiological phenotypes. The tolerant ecotype (CC1009) successfully adapted, developing a functional CCM with a structured starch sheath. Conversely, the sensitive ecotype (CC2343) suffered growth arrest and formed malformed pyrenoids. Transcriptomics revealed that CC1009 initiated a rapid initial response, upregulating chloroplast proteostasis and downregulating nucleotide metabolism. CC2343 showed a massive, delayed transcriptional response, downregulating genes coding for photosystems and tetrapyrrole biosynthesis. This unbiased transcriptomic approach identifies key candidate genes driving algal acclimation to hyperoxic stress in natural, high-light environments.

## INTRODUCTION

Green algae are highly successful primary producers in dynamic environments, including lakes and soil, where they face selective pressure from fluctuating temperature, light quality, and nutrient availability. Unsurprisingly, algae have evolved highly flexible metabolic systems that allow them to maintain photosynthesis despite sudden environmental changes. One classic example is the flexible deployment of carbon-concentrating mechanisms (CCMs) that allow phototrophic growth even under low CO_2_ availability (Giordano et al., 2005). Due to relatively slow gas diffusion rates in aquatic environments, algae can experience significant daily fluctuations of dissolved CO_2_ and O_2_. This is especially true for highly photosynthetically active algal cultures exposed to direct sunlight during the day, which produce oxygen endogenously at high rates (Peng et al., 2013). At night, photosynthesis is inactive and O_2_ consumption by respiration results in anoxia (Quinn et al., 2000; Quinn et al., 2002; Hemschemeier et al., 2013). Conversely, during the day, photosynthetic water oxidation generates O_2_ that accumulates in the surrounding environment reaching supersaturating (i.e., hyperoxic) levels up to four times higher than air saturation (Peng et al., 2013). Hyperoxia can lead to decreased photosynthetic efficiency, energy loss via photorespiration and oxidative damage to cellular components (Marquez et al., 1995; Ugwu et al., 2007). Additionally, the depletion of available CO_2_ through assimilation further elevates the O_2_:CO_2_ ratio, exacerbating the risk of O_2_ incorporation and photorespiration.

One major consequence of photosynthesis under these conditions is the accelerated production of reactive oxygen species (ROS), which can damage DNA, lipids, and proteins (Apel and Hirt, 2004). In chloroplasts, photochemistry yields unstable, high-energy intermediates that interact with molecular O_2_ to generate ROS. Singlet oxygen (^1^O_2_) is continuously created in photosystem II (PSII) in the light, resulting in the formation of triplet chlorophyll and subsequent energy transfer to molecular O_2_ (Durrant et al., 1990; Hideg et al., 1994). Concurrently, the photoreduction of O_2_ yields superoxide (O_2_^-^), which superoxide dismutase enzymatically converts to hydrogen peroxide (H_2_O_2_) (Asada, 1999). While excessive ROS is toxic, these molecules also serve as crucial retrograde signals regulating nuclear gene expression (Fischer et al., 2005; Wakao et al., 2014; Blaby et al., 2015; Ma et al., 2020). However, despite ROS being a direct byproduct of photosynthesis, comprehensive studies in phototrophically grown algae are notably lacking. This is a critical omission, as a reliance on photosynthesis for growth fundamentally alters the cellular energetic and redox landscape, thereby changing how cells perceive and respond to oxygen.

Instead, current research relies primarily on the exogenous chemical application of H_2_O_2_ (Blaby et al., 2015) and ^1^O_2_-producing photosensitizing dyes (Fischer et al., 2005; Wakao et al., 2014; Ma et al., 2020; Pancheri et al., 2024) in photoheterotrophically grown *Chlamydomonas reinhardtii* (hereafter Chlamydomonas). These chemical treatments confound gene expression analysis because extracellular ROS must diffuse across multiple membranes (e.g., plasma membrane, chloroplast envelope, mitochondrial membranes) to reach target organelles, each equipped with its own distinct signaling and detoxification networks. Because different ROS species possess varying membrane permeabilities and half-lives, organellar concentrations cannot be precisely controlled. Crucially, photoheterotrophic growth suppresses native chloroplast development and pyrenoid dynamics, making it difficult to distinguish true *in vivo* physiological adaptations from generic, non-specific stress responses.

To circumvent these limitations, we exploited the natural genetic variation of Chlamydomonas using two strains with vastly contrasting phenotypes under hyperoxia: the tolerant ecotype CC1009 and the sensitive ecotype CC2343 (Neofotis et al., 2021; Lucker et al., 2022).

Phenotypic observations reveal that under hyperoxia, CC1009 accumulates intracellular H_2_O_2_ while successfully maintaining a functional CCM and a well-structured pyrenoid. In contrast, CC2343 fails to elevate intracellular H_2_O_2_, exhibits malformed pyrenoids, and undergoes rapid growth arrest. Based on these divergent phenotypes, we hypothesized that hyperoxia functions as a distinct physiological signal, potentially mediated by intracellular H_2_O_2_, that triggers a coordinated restructuring of photosynthetic components to activate the CCM, initiate pyrenoid formation and photoprotection. Alternatively, hyperoxia may act merely as destructive oxidative stress, causing a systemic homeostatic collapse in the sensitive genetic background rather than an adaptive response.

Here, we tested these competing hypotheses by analyzing the genome-wide transcriptional responses of CC1009 and CC2343 to hyperoxia under strict phototrophic conditions using RNA-seq. By integrating functional enrichment (KEGG, GO, and MapMan pathways) with comparative analyses against published low-CO_2_ and exogenous ROS datasets, we map the core regulatory networks governing the algal response to high O_2_:CO_2_ ratios. Ultimately, our transcriptomic profiling demonstrates that tolerance is driven by a highly targeted, adaptive signaling program that maintains photosynthetic performance, whereas sensitivity reflects a generalized failure of cellular homeostasis under oxidative stress, culminating in photosynthetic collapse.

## RESULTS

### Chlamydomonas ecotypes CC1009 and CC2343 are genetically diverse

We previously identified striking phenotypic differences between Chlamydomonas ecotypes CC1009 and CC2343 regarding their capacity to adapt to hyperoxia (Neofotis et al., 2021; Lucker et al., 2022). While CC1009 maintains phototrophic growth after the onset of hyperoxia, CC2343 enters growth arrest within the first 10h after exposure. We reproduced this growth difference here, it is clearly demonstrated by the much greater dilution factors required for CC1009 during turbidostatic cultivation in the environmental photobioreactors (ePBRs) (Figure S1). Both ecotypes adapt to hyperoxia by inducing pyrenoids, structural components of the CCM, albeit with distinct differences in the physical appearance of the starch sheath. Under hypoxic conditions, the CC1009 pyrenoid starch sheath is robust and contiguous, whereas the CC2343 starch sheath appears underdeveloped and disjointed. Additionally, CC2343 exhibits lower productivity, reduced rubisco activity, and lower O_2_ compensation points under hyperoxia (Neofotis et al., 2021).

Although previous studies provided significant genomic information on previous Chlamydomonas ecotypes (Flowers et al., 2015; Gallaher et al., 2015), it remained unclear to what extend genetic variation may explain observed phenotypic differences between the two ecotypes. To determine the genomic differences between the two strains, we reanalyzed previously sequenced genomic DNA from both ecotypes (Lucker et al., 2022). We found that CC1009 shares 99.52% sequence identity with the reference genome CC4532, which is notably higher than the 95.81% identity it shares with CC2343. Meanwhile, CC2343 shares 95.93% sequence identity with the CC4532 reference (Table S1). When compared to CC4532, strain CC2343 carries approximately nine times more single-nucleotide polymorphisms (519,400 versus 74,622) and about four and a half times as many insertions or deletions (14,900 versus 3,285) than strain CC1009. No translocations or inversions were identified. Studies investigating various aspects of Chlamydomonas metabolism suggest phenotypic variation among wild-type laboratory strains. This variation typically is attributed to the presence of two genetic haplotypes distributed in a mosaic pattern across these strains (Glaesener et al., 2013; Gallaher et al., 2015).

### Distinct transcriptional responses in two ecotypes exposed to hyperoxia

We surveyed the genome-wide transcriptional response of CC1009 and CC2343 to hyperoxia using RNA-seq. Cells were grown photoautotrophically in ePBRs under high light (2000 µmol photons m^-2^s^-1^ PFD) and hyperoxia (5% CO_2_, 95% O_2_), mimicking a bright, sunny day in nature. Samples were collected at 0 h baseline (5% CO_2_, 21% O_2_), 1, 3 and 6 h after the onset of hyperoxia to capture immediate adaptation responses, as well as 24 and 48 h to monitor long-term adaptation strategies. While alignment to the reference genome yielded consistent mapping efficiencies across all timepoints, CC2343 exhibited a lower unique read mapping percentage (79.5%) compared to CC1009 (91.7%) (Figure S2), while independent time points showed excellent correlation (Figure S3). The observed difference in unique read mapping percentage is primarily attributed to a higher proportion of unmappable short reads in CC2343 (14.4%) relative to CC1009 (3.8%) (Figure S2). To parse this rich dataset, we performed a principal component analysis (PCA) to reduce complexity (Figure 1A). The strong differences in hyperoxia adaptation between the two Chlamydomonas strains are clearly illustrated by their separation along Principal Component 1 (PC1, Figure 1A), which captures over 94% of the total variance and separates the samples by ecotype (Figure 1A). Conversely, Principal Component 2 (PC2) sorts the samples chronologically by hyperoxia exposure duration. Both ecotypes display a similar upward trajectory along this axis (Figure 1A). Visualizing these temporal transcriptomic shifts underscores the extent to which individual gene expression profiles are driven at a genome-wide level.

**Figure 1.**
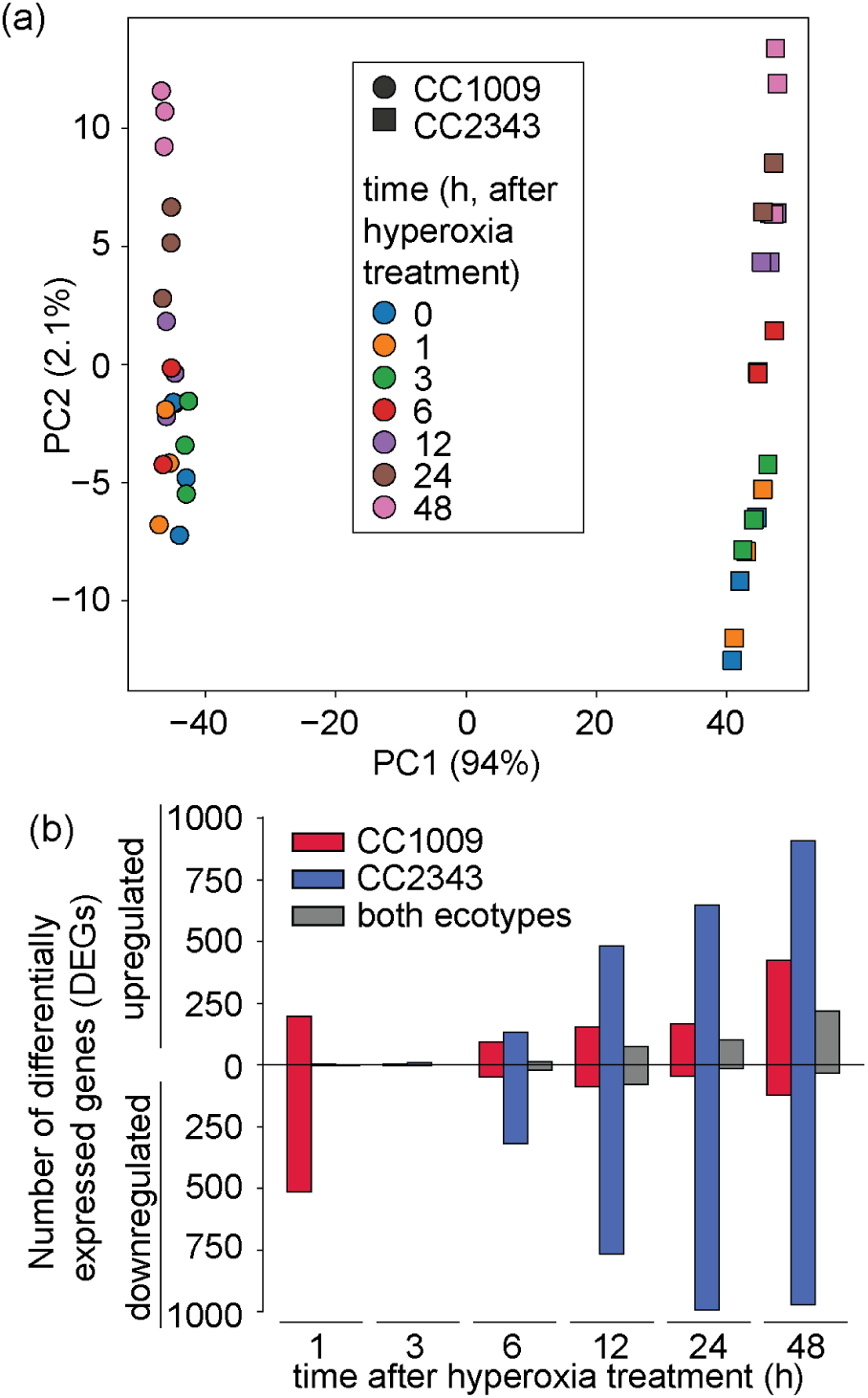
Chlamydomonas ecotypes CC1009 and CC2343 exhibit distinct, transcriptional responses to hyperoxia. (a) Principal component analysis. 2D PCA plot showing sample distribution along principal component 1 (PC1) and principal component 2 (PC2). Parentheses represent percentage variance explained by the component. Ellipses group datapoints by ecotype. (b) Shown is the number of significant differentially expressed genes (DEGs) unique to CC1009 (red), unique to CC2343 (blue), or present in both ecotype (gray). Upregulated and downregulated genes are shown as indicated. DEGs are defined genes with a LFC > 1 and an adjusted p-value < 0.05.

The two ecotypes exhibited distinct molecular responses to hyperoxia treatment, varying significantly in both timing and amplitude of transcriptional changes (Data S1). Within the first hour of exposure, the expression of hundreds of transcripts was altered in CC1009 (199 up and 515 down, Figure 2A), whereas only nine total transcripts were differentially expressed in CC2343 at the same timepoint (Figure 1B). Conversely, subsequent changes are much more pronounced in the sensitive strain (CC2343, Figure 1B). Given that CC1009 is hyperoxia-tolerant, we hypothesized that its rapid transcriptional reprogramming represents a crucial strategy for survival and adaptation under oxidative stress, whereas the delayed response in CC2343 reflects a failure to acclimate. To identify functional pathways critical to tolerating oxidative stress, we analyzed each set of differentially expressed genes (DEGs) for functional pathway enrichment using the Algal Functional Annotation Tool (Lopez et al., 2011). Gene ontology (GO) terms (Ashburner et al., 2000; Aleksander et al., 2023), KEGG pathways (Kanehisa and Goto, 2000), and MapMan terms (Thimm et al., 2004) were evaluated for each ecotype across all timepoints (Figure 2 and Data S2).

**Figure 2.**
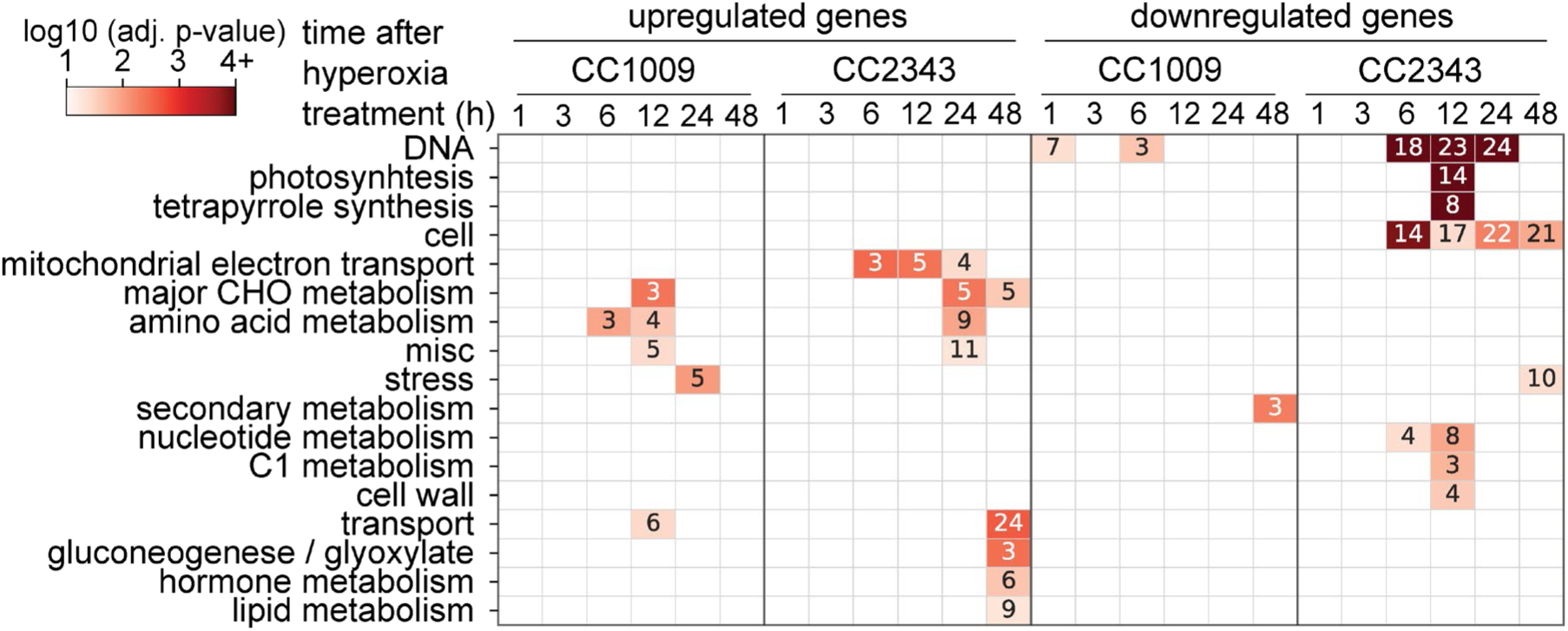
CC2343 downregulates transcription of genes encoding proteins involved in DNA metabolism and photosynthesis in response to hyperoxia. Functional enrichment of MapMan level 1 terms. The values in each cell represents the number of hits for each enriched term. The color bar scale represents the -log10 p-value from 1 (light red) to greater than 4 (dark red) for each enriched term. Terms with less than 2 hits or a p-value > 0.05 were omitted.

Surprisingly, despite the maintenance of phototrophic growth, a subset of genes rapidly downregulated in CC1009 within the first hour of hyperoxia encode proteins involved in nucleic acid metabolism (Figures 1b and 2, Data S1). Notable examples include RNA metabolism genes such as exoribonuclease (*XRN2*, Cre03.g152100), snRNA-associated Sm-like small nuclear riboproteins (*SMP6B,* Cre06.g251300; Cre10.g441950) and DEAD/DEAH box helicases (Cre06.g282600, Cre16.g662000). Genes involved in DNA replication and processing were similarly repressed, including subunits of the DNA-melting helicase such as minichromosome maintenance proteins (*MCM4*, Cre07.g316850; *MCM5*, Cre01.g023150) and alpha subunits of the DNA polymerase (*POLA2*, Cre01.g017450). The latter function in DNA replication and are typically co-expressed with histone genes during DNA replication and cell division (Strenkert et al., 2019).

In contrast, genes coding for proteins involved in protein homeostasis and high-light stress acclimation were upregulated in the tolerant CC1009 strain. These include the ClpB chaperone (*CLPB3*), Chaperonin 10 (*CPN10*), Plastid terminal oxidase 1 (*PTOX1*), Vesicle-inducing protein in plastids 2 (*VIPP2*) and Early light-induced LHC-like protein (*ELIP7*). Interestingly, although *VIPP2* was significantly upregulated in CC1009, it maintained a higher absolute baseline expression in CC2343 (26 mean FPKM in CC1009 vs. 231 mean FPKM in CC2343). VIPP2 localizes to the thylakoid membrane, is induced by high light and H_2_O_2_ stress (Theis et al., 2020) and is proposed to act as a chloroplast membrane stress sensor that interacts with heat shock protein HSP22E/F to alleviate membrane stress (Theis et al., 2020). Corroborating this, *HSP22E/F* expression was significantly upregulated in both ecotypes. Additionally, the chloroplast *CLPB3* gene, which colocalizes with *HSP22E/F* during H_2_O_2_-producing heat stress (Kreis et al., 2023), was also upregulated in both strains.

These data demonstrate that CC1009 initiates a rapid transcriptional defense mechanism against hyperoxia that is absent in CC2343. The hyperoxia-sensitive CC2343 cells delayed transcriptomic modulation until approximately six hours post-treatment (Figure 1B).

Furthermore, functional enrichment analyses revealed that the DEGs in CC2343 targeted entirely different biological processes (Figure 2), culminating in a severe, immediate negative impact on photosynthesis. Specifically, at six hours after the onset of hyperoxia, CC2343 strongly downregulated MapMan Level 1 categories associated with DNA, photosynthesis, tetrapyrrole synthesis, cell, stress, nucleotide metabolism, C1-metabolism, and cell wall (Figure 2). This dramatic collapse in the photosynthetic machinery of CC2343 included the downregulation of photosystem 1 (PSI) components (*PSAD*, *PSAF*, *PSAG*, *PSAH*, *PSAI*, *PSAK*, *PSAL*, *PSAN*, *PSAO*), PSII components (*PSBO*, *PSBP1*, *PSBQ*, *PSBR*, *PSBX*, *PSBY2*), the cytochrome *b*6*f* complex (*PETM*, *PETN*, *PETO*), PSI light harvesting complexes (*LHCA2*, *LHCA3*, *LHCA4*, *LHCA5*, *LHCA6*, *LHCA7*, *LHCA8*), PSII light harvesting complexes (*LHCBM2*, *LHCBM3*, *LHCBM4*, *LHCBM5*, *LHCBM6*, *LHCBM7*, *LHCBM8*), and chloroplast ATP synthase subunits (*ATPC*, *ATPD*, *ATPG*) (Figure 3). A similar trend was observed for tetrapyrrole biosynthesis, where transcripts encoding chlorophyll synthetase (*CHLG1*), magnesium chelatase subunits (*CHLI1*, *CHLD1*), Chlorophyllide a oxygenase (*CAO1*), and protochlorophyllide reductase (*POR1*) were significantly depleted 12 hours post-exposure (Data S1 and Data S2). Interestingly, this trend was also observed in the tolerant CC1009 strain, but only as a delayed response during long-term hyperoxia treatment between 12 and 48 h (Figure 3).

**Figure 3.**
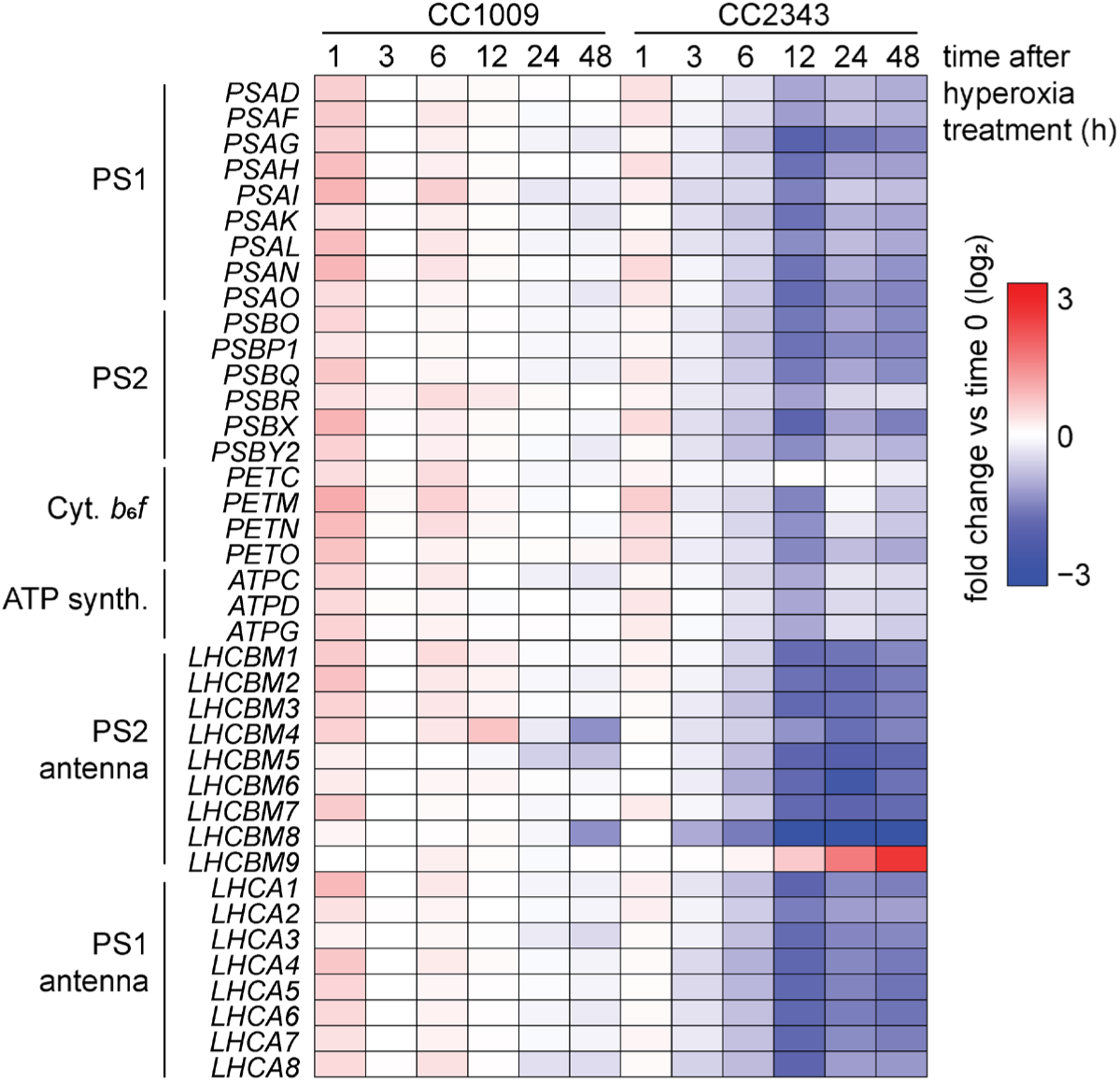
Transcript abundances of genes encoding proteins involved in photosynthesis are downregulated in CC2343 in response to hyperoxia. Shown are relative RNA abundance data of genes encoding subunits of the photosynthetic electron transfer chain in CC1009 and CC2343. Shown are averages of three independent replicates.

### Ecotype-specific induction of *LCI* and CCM-related genes

The Chlamydomonas carbon concentrating mechanism (CCM) centralizes CO_2_ near rubisco within a specialized starch-sheathed, phase separated organelle called the pyrenoid via a vast spatial network of proteins (Mackinder et al., 2017). CO_2_ limitation (∼0.04% CO_2_) promotes pyrenoid assembly and induces extensive changes at the transcript level, notably upregulating carbonic anhydrases, bicarbonate transporters, and CO_2_ transporters (Brueggeman et al., 2012; Fang et al., 2012), which collectively optimize CO_2_ fixation rates while suppressing photorespiration. Many of these low CO_2_-responsive genes are under the control of the master regulator cia5 (Xiang et al., 2001; Wang et al., 2005; Fang et al., 2012), yet the precise mechanism governing CCM activation remains unknown. Work by Santhanagopalan and colleagues suggests that a signal derived from photorespiration or the redox state of the photosynthetic electron transport (PET) chain may trigger CCM induction (Santhanagopalan et al., 2021). Corroborating this, our previous work demonstrated that hyperoxia, a condition that accelerates photorespiration, stimulates reactive oxygen species (ROS) generation, and reduces PET efficiency, induces pyrenoid formation and activates the CCM, even under elevated (5%) CO_2_ condition (Neofotis et al., 2021). However, while hyperoxia drives functional CCM induction in CC1009, it results in atypical, malformed starch sheaths in the CC2343 ecotype. These morphological differences raise a fundamental question: is the transcriptional response of core CCM components comparable or distinct between the two genetically diverse ecotypes?

We compared hyperoxia-induced transcriptomic changes in our datasets within the canonical carbon-limitation response from previous work (Fang et al., 2012). After mapping reads to the Chlamydomonas reference genome v6.1 (Craig et al., 2023), we systematically defined the core low CO_2_ inducible (*LCI*) regulon by evaluating transcriptional responses under low (L) and very low (VL) CO_2_ conditions relative to ambient controls (Data S3). For upregulated genes, we identified 68 transcripts in the L condition, 250 transcripts in the VL condition, and an overlap of 413 genes induced by both treatments. Conversely, downregulation affected 86 genes in L, 468 in VL, and 359 shared across both conditions.

Based on these stringent comparative criteria, we established a final reference gene set comprising 731 upregulated and 913 downregulated canonical *LCI* genes to cross-reference against our ecotype-specific hyperoxia datasets.

We found pronounced differences in the expression of many *LCI* genes between CC1009 and CC2343 (Figure 4A). In general, more CCM-related genes were upregulated in CC1009 in response to hyperoxia, particularly at early time points, whereas more genes were downregulated in CC2343. Among the most divergent were *LCI9* and *SSS2*, both of which encode proteins involved in pyrenoid starch metabolism.

**Figure 4.**
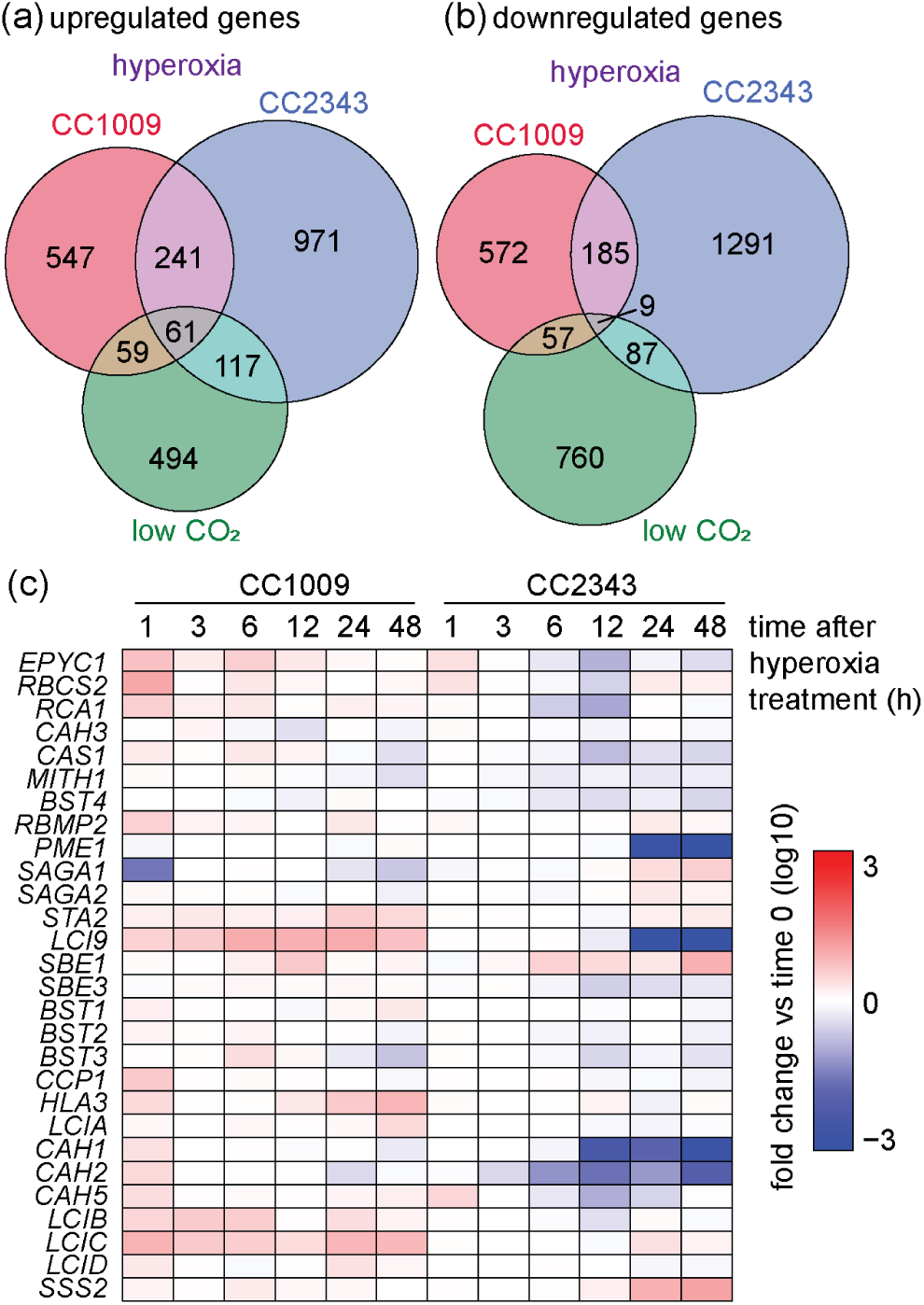
Commonalities between hyperoxia and the low CO_2_ response in CC1009 and CC2343. (a) Shown are overlapping and distinctly differentially expressed genes in response to cell grown under hyperoxia (this study) as compared to low CO_2_ grow cells (data from Brueggeman et al.) (c) Shown are relative RNA abundance data of selected genes encoding low CO_2_ inducible genes. Shown are averages of three independent replicates.

*LCI9* transcripts were abundant in CC1009 (mean FPKM 28) but nearly absent in CC2343 (mean FPKM 3; Figure 4). LCI9 forms a mesh-like network linking individual starch plates within the pyrenoid matrix and contains carbohydrate-binding domains that may mediate starch remodeling (Mackinder et al., 2017). Consistent with this role, LCI9-deficient mutants fail to assemble the pyrenoid starch sheath properly and exhibit severe growth defects under CO₂- limiting conditions (Adler et al., 2025).

Similarly, *SSS2*, which encodes soluble starch synthase II and contributes to pyrenoid starch synthesis following cell and pyrenoid division (Fletcher, 2020), was expressed at much higher levels in CC1009 than in CC2343 (mean FPKM 187 vs. 23). The gene encoding Pyrenoid Membrane Enriched 1 (*PME1*), a protein that spans the pyrenoid matrix, also showed markedly higher expression in CC1009 (mean FPKM 100 vs. 3). Although PME1 is not essential for pyrenoid assembly or growth under low CO₂ (Franklin et al., 2025), the coordinated divergence in expression of *LCI9*, *SSS2*, and *PME1* suggests that their regulation may be important for pyrenoid structural organization and proper alignment of starch plates under hyperoxia.

### Comparison of RNA expression responses to oxidative stress treatments

Hydrogen peroxide (H_2_O_2_) is produced as a natural consequence of oxygenic photosynthesis, particularly under high light, carbon-limiting environments (Roach et al., 2015). High O_2_ and even low levels of exogenous H_2_O_2_ levels have recently been shown to strongly induce pyrenoid formation and activate the CCM, even in the presence of elevated CO_2_ (Neofotis et al., 2021).

Interestingly, the hyperoxia-tolerant strain CC1009 increases intracellular H_2_O_2_ concentrations under hyperoxia, whereas H_2_O_2_ levels in the hyperoxia-sensitive strain CC2343 remain constant (Neofotis et al., 2021). While H_2_O_2_ serves as a critical signaling molecule, other reactive oxygen species (ROS), such as singlet oxygen (^1^O_2_), are also generated during photosynthesis and can trigger broad stress responses. To distinguish between a specific hyperoxia-induced signaling pathway and a generalized response to singlet oxygen, we compared our results with publicly available datasets investigating the effects of Rose Bengal (RB) and high light (HL) (Pancheri et al., 2024).

To minimize the confounding effects of exogeneous ^1^O_2_ production by RB, we defined a core ^1^O_2_-responsive gene set containing only those DEGs common to both the RB and HL treatments (Data S3). A comparison of upregulated genes yielded 652 overlapping DEGs, with 3,845 genes specific to the RB treatment (out of 4,497 total) and 945 specific to HL (out of 1,597 total). Conversely, analysis of downregulated genes yielded 1,414 intersecting DEGs, with 4,722 genes specifically repressed by RB (out of 6,136 total) and 535 specifically repressed by HL (out of 1,949 total).

The core set of 652 upregulated ^1^O_2_-responsive genes was compared against upregulated gene sets from ecotypes CC1009 (908 genes) and CC2343 (1,390 genes), identifying 57 genes that were consistently upregulated across all three datasets. Pairwise comparisons revealed 245 genes shared exclusively between CC1009 and CC2343, 127 genes shared between CC2343 and ^1^O_2_, and 38 genes shared between CC1009 and ^1^O_2_ (Figure 5A). Notably, the hyperoxia sensitive strain CC2343 shared more than three times as many upregulated genes with the core ^1^O_2_ stress dataset than the tolerant strain CC1009, providing initial evidence that hyperoxia triggers a more severe, generalized oxidative stress state in the sensitive background.

**Figure 5.**
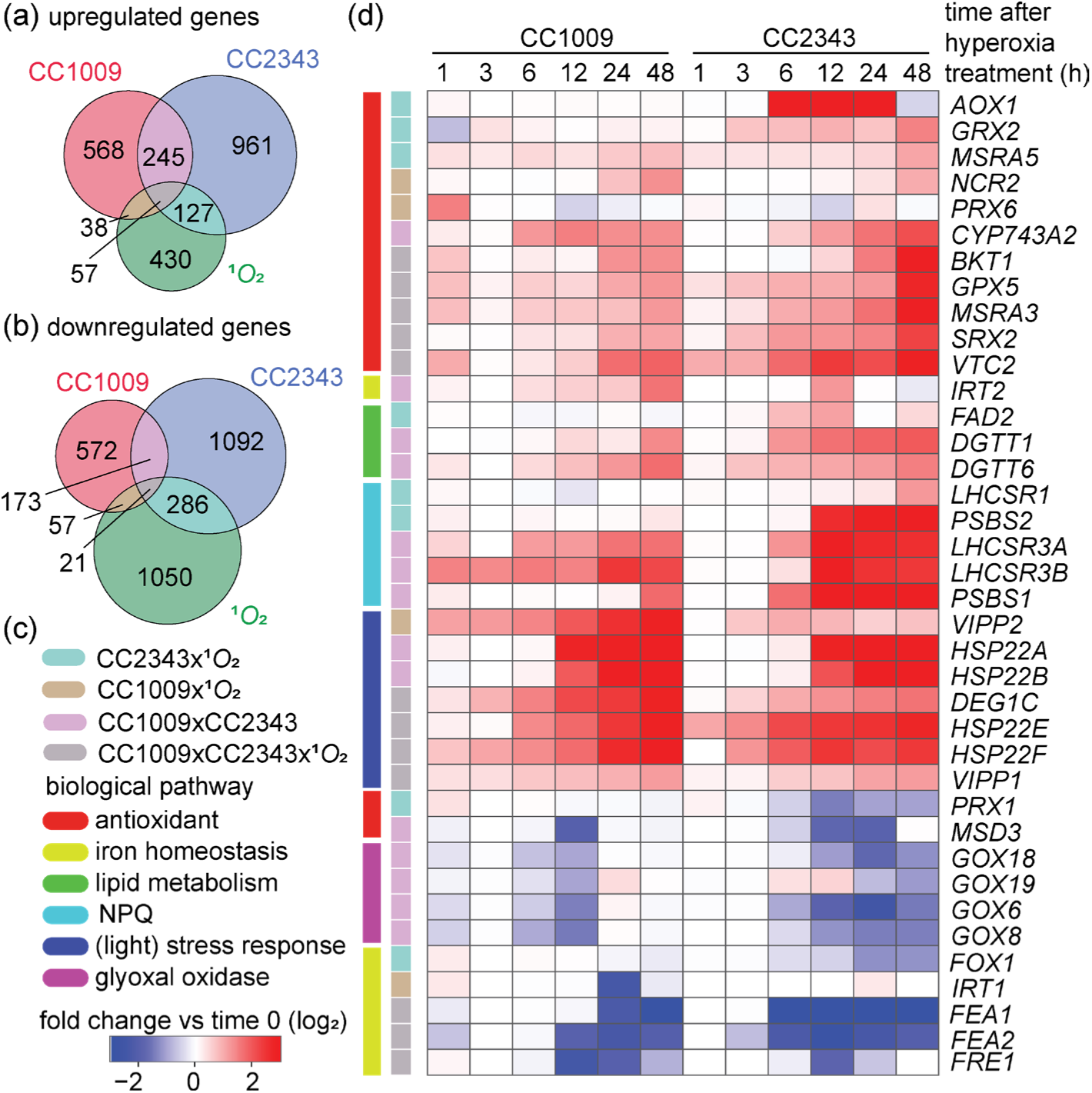
Commonalities and difference between gene expression profiles of CC1009 and CC2343 compared to single oxygen responses. (a) Shown are overlapping and distinctly differentially expressed genes in response to cell grown under hyperoxia (this study) as compared to singlet oxygen inducing conditions (highlight and rose Bengal treatment, data from Roach et al.) (c,d) Shown are relative RNA abundance data of selected genes encoding low ^1^O_2_ inducible genes. Shown are averages of three independent replicates.

Parallel analyses of 1,414 core downregulated ^1^O_2_ genes against the downregulated genes of CC1009 (823 genes) and CC2343 (1,572 genes) revealed 21 DEGs shared among all three conditions (Figure 5B). Pairwise comparisons showed 173 DEGs shared between CC1009 and CC2343, 286 shared between CC2343 and ^1^O_2_, and 57 shared between CC1009 and ^1^O_2_ set (Figure 5).

Several antioxidant and photoprotective genes were consistently upregulated across all three datasets, including Beta-carotene ketolase (*BTK1*), methionine-S-sulfoxide reductase (*MSRA3*), Glutathione peroxidase 5 (*GPX5*), sulfiredoxin (*SRX5*), and the GDP-L-galactose phosphorylase (VTC2). This core stress regulon also included the Deg protease (*DEG1C*), Vesicle inducing protein in plastids 1 (*VIPP1*) and Heat shock protein 22E/F (*HSP22E/F*). In contrast, a common set of downregulated genes was enriched for proteins involved in iron metabolism, such as Ferric-chelate reductase/oxidoreductase (*FRE1*) and Fe-assimilation protein (*FEA1, FEA2*).

Transcripts upregulated in both hyperoxia datasets (CC1009 and CC2343) but not differentially expressed by ^1^O_2_ included *PSBS1*, cysteine dioxygenase (*CDO2*), heat shock protein 22A/B (*HSP22A/B*), cytochrome P450 (*CYP743A2*), diacylglycerol acyltransferase (*DGTT6*), and the iron-nutrition transporter *IRT2*. Conversely, co-downregulated genes in both ecotypes included glyoxal oxidases (*GXO6/8/18/19*), light-harvesting chlorophyll *a*/*b*-binding proteins (*LHCBM4/8*), manganese superoxide dismutase (*MSD3*), and the light-harvesting chlorophyll a/b stress-related protein 3 (LHCSR3). Notably, LHCSR3, encoded by two identical genes (*LHSCR3A* and *LHCSR3B*) codes for a protein essential for qE-type non-photochemical quenching (NPQ) of excess light in Chlamydomonas (Peers et al., 2009; Tokutsu and Minagawa, 2013). Both genes exhibited upregulation exclusively under hyperoxia, consistent with previous research (Roach et al., 2020).

Evaluating ecotype-specific overlaps with the ^1^O_2_ dataset highlighted divergent acclimation strategies. The hyperoxia-tolerant CC1009 and the ^1^O_2_ datasets shared the upregulation of peroxiredoxin Q (*PRX6*) and vesicle-inducing protein in plastids 2 (*VIPP2*), while concurrently downregulating the iron transporter *IRT1* and the voltage-dependent anion-selective channel protein 2 (*VDAC2*).

In contrast, the hyperoxia-sensitive CC2343 strain and the ^1^O_2_ dataset shared an extensive list of upregulated photoprotective and stress-responsive genes, including *PSBS2*, Deg protease *DEG1A*, alternative oxidase 2 (*AOX2*), early light-induced LHC-like proteins (*ELIP4/10*), light-harvesting chlorophyll stress-related protein 1 (*LHCSR1*), methionine-S-sulfoxide reductase (*MSRA5*), glutaredoxin 2 (*GRX2*), fatty acid desaturase 2 (*FAD2*), methionine sulfoxide reductase 1B (*MSR1B*), and glutathione S-transferase (*GSTS1*). Downregulated genes common to CC2343 and ^1^O_2_ included ferredoxin 2 (*FDX2*), 2-Cys peroxiredoxin (*PRX1*), multicopper ferroxidase (*FOX1*), acetate transporters (*GFY3/4/5*), and light-harvesting chlorophyll *a*/*b*-binding proteins (*LHCBM6/7/15*).

## DISCUSSION

### How supersaturating levels of oxygen affect algal metabolism – a sunny day in the life of a green alga

It has been long been proposed that *Chlamydomonas reinhardtii* induces its carbon-concentrating mechanism (CCM) as a specific response to low CO_2_ availability. The current data challenges this framework by demonstrating that hyperoxia, even in the presence of high CO_2_, can also activate the CCM. Hyperoxia is ecologically highly relevant because O_2_ levels can reach extremely high levels during peak photosynthesis in dense algal mats, competing with CO_2_ at rubisco, diverting photosynthetic fluxes to energy-losing photorespiration.

Our comparative transcriptomic analysis suggests that the capacity to survive hyperoxic conditions relies on a rapid, highly coordinated reprogramming of the cellular transcriptome. In the tolerant ecotype CC1009, this is marked by an immediate suppression of nucleic acid metabolism and cell division machinery within the first hour, effectively pausing the cell cycle to redirect metabolic resources toward stress mitigation. Concurrently, CC1009 selectively upregulates critical chaperones and potential membrane stabilizers, such as *VIPP2* and *CLPB3*. Crucially, our findings illuminate a signaling paradigm regarding intracellular hydrogen peroxide (H_2_O_2_). While excess reactive oxygen species (ROS) are inherently damaging, the transient accumulation of intracellular H_2_O_2_ observed in CC1009, but not in CC2343, (Neofotis et al., 2021) likely functions as a signal required to orchestrate this rapid defense. Conversely, the hyperoxia-sensitive strain CC2343 fails to elevate its intracellular H_2_O_2_ pool upon exposure, resulting in a disastrous 6-hour delay in transcriptomic modulation. When CC2343 finally responds, it does not execute a protective acclimation program; instead, it undergoes a systemic collapse of its entire photosynthetic apparatus, characterized by the profound downregulation of core photosynthetic complexes such as PS1, PS2, cytochrome *b*_6_*f*. This transcriptional pattern indicates that a failure to perceive or propagate early oxidative signals prevents the activation of essential photoprotective mechanisms, leading to irreversible growth arrest and cell death.

### Natural genetic diversity dissects genotype-to-phenotype connections

Collectively, these results demonstrate that the phenotypic divergence between Chlamydomonas ecotypes CC1009 and CC2343 under hyperoxia is the product of profound underlying genomic variation. Previous research established that common Chlamydomonas laboratory strains and field isolates exhibit significant genetic and phenotypic diversity (Flowers et al., 2015; Gallaher et al., 2015). Our findings support this observation: the substantial differences in SNPs and indels between CC1009 and CC2343 relative to the reference genome highlight a deep evolutionary divergence, suggesting that these two ecotypes have been shaped by distinct selective pressures in their native environments. For instance, the lake environment from which CC1009 was isolated may have exerted selective pressure for robust pyrenoid formation due to temperature variation, low CO_2_ availability, and O_2_ fluctuations. In contrast, the soil environment of CC2343, characterized by different gas exchange dynamics and high heavy-metal content, may have favored an entirely different set of metabolic priorities. These divergent evolutionary histories likely underpin the distinct regulatory capacities we observed in these ecotypes regarding carbon assimilation and pyrenoid formation under hyperoxic stress

This genotype-to-phenotype connection is clearly illustrated by the failure of CC2343 to induce core low-CO_2_-inducible (*LCI*) genes encoding structural components of the CCM. The assembly of a functional starch sheath around the pyrenoid matrix is critical to prevent CO_2_ leakage and protect rubisco from oxygenation. Our data suggest that the underdeveloped and disjointed starch sheath phenotype of CC2343 under hyperoxia correlates with its inability to induce *LCI9* and *SSS2*, which are essential for structural plate alignment and starch biosynthesis, respectively (Mackinder et al., 2017; Adler et al., 2025).

Because the upstream transcription factor CIA5 is present and seems structurally intact in CC2343, this ecotype-specific silencing of *LCI9* and *SSS2* suggests that the cis-regulatory elements or secondary sensor networks required to relay the oxidative signal to these specific promoters are defective or absent in the CC2343 genetic background. Consequently, utilizing such natural ecotypic variation provides a powerful genetic framework to isolate the polymorphism networks that dictate the structural fidelity of the CCM and overall cellular robustness against environmental oxidative stress.

### Hyperoxia-induced photoprotection and the role of non-photochemical quenching (NPQ)

Our findings highlight the critical role of non-photochemical quenching (NPQ) as a rapid energy dissipation mechanism, which is essential in the oxygenated environments characteristic of actively photosynthesizing green algal cultures. A key gene coding for the qE-Type NPQ protein, LHCSR3, is significantly upregulated across both ecotypes in response to hyperoxia.

This robust induction suggests an active, fundamental defense mechanism against high endogenous oxygen levels, consistent with previous observations regarding the induction of photoprotective mechanisms during oxidative stress (Wakao et al., 2014; Blaby et al., 2015; Roach et al., 2020; Pancheri et al., 2024).

However, the temporal dynamics of this photoprotective response reveal divergent acclimation strategies between the two ecotypes. While both strains attempt to manage excess excitation energy, the hyperoxia- sensitive strain CC2343 displays a delayed, compensatory induction of auxiliary NPQ components. Specifically, both *PSBS1* and *PSBS2* showed a 5-fold increase in expression after 48 hours of hyperoxia exposure in CC2343. In Chlamydomonas, PSBS acts transiently to facilitate rapid NPQ activation, typically operating in tandem with, or in partial compensation for, LHCSR3-dependent NPQ pathways (Peers et al., 2009; Tibiletti et al., 2016; Strenkert et al., 2019). The late, pronounced upregulation of *PSBS1* and *PSBS2* in CC2343 likely represents a late, molecular compensatory mechanism to dissipate persistent excitation energy and mitigate the effects of photoinhibition resulting from the collapse of its photosynthetic apparatus.

### The molecular specificity of hyperoxia versus generalized ROS damage

By cross-referencing transcriptome data with transcriptomes form algal cells exposed to high light or Rose Bengal (Pancheri et al., 2024), we delineated the molecular signature of hyperoxia from canonical, chemically-induced singlet oxygen (^1^O_2_) stress responses. The shared induction of a core regulon across all datasets suggests that hyperoxia acclimation shares fundamental, molecular responses with standard, oxidative stress pathways. Alternatively, differences in gene expression may be attributed to the phototrophic, high-light conditions which differ significantly from the photoheterotrophic, low-light conditions employed by Pancheri. Nevertheless, the distinct partitioning of the remaining molecular changes underscores how *in vivo*, endogenous oxygen supersaturation differs from exogenous chemical induction of ^1^O_2_.

We noticed that the hyperoxia-tolerant ecotype CC1009 coordinates its response to hyperoxia with the transcriptional upregulation of genes known to stabilize thylakoid membranes during oxidative stress like *VIPP2* and peroxiredoxins such as *PRX6* (Dayer et al., 2008; Theis et al., 2020), while avoiding a more widespread, transcriptional response seen in cells after chemical treatments. In stark contrast, the sensitive strain CC2343 exhibits an extensive transcriptional overlap with the ^1^O_2_ dataset, inducing a wide array of stress-responsive genes such as *AOX2*, *ELIP4/10*. This indicates that, while CC1009 induces a highly specific transcriptional response to maintain chloroplast homeostasis, the genetic lesions in CC2343 result in an inability to adapt to hyperoxia and lead to a more generalized, untargeted transcriptional response.

Our comparative analysis demonstrated that hyperoxia represents a physiologically distinct stress condition that cannot be adequately mimicked using exogenous chemical treatments such as Rose Bengal. While these chemical treatments result in a sudden, unlocalized ROS accumulation, naturally occurring hyperoxia drives a localized, specific signaling cascade. For tolerant ecotypes, this endogenous ROS signal is successfully integrated into a protective, developmental program that activates the CCM and reorganizes chloroplast architecture.

## DATA AVAILABILITY

Raw sequence data are available in the NCBI Sequence Read Archive (BioProject PRJNA1477029) and will be made publicly available after publication. The analysis pipeline, including Python scripts and Jupyter notebooks, is maintained in a public GitHub repository (https://github.com/templj/hyperoxia-rna).

## ACKNOWLEDGEMENTS

This material is based upon work supported primarily by the U.S. Department of Energy, Office of Science, Office of Basic Energy Sciences, United States Department of Energy under Award Number DE-FG02-91ER20021. The work was supported by the National Science Foundation Research Traineeship Program (DGE-1828149, to Joshua A. Temple), ExxonMobil Research and Engineering Company and the Jan IngenHousz Institute (to DMK). The development of the methods and instruments used in this work were supported by the U.S. Department of Energy Office of Biomass Program grant (DE-EE0003046). This work was supported by grants DE- SC0007101 and DE-FG02-09ER20310 from the Photosynthetic Systems program from Division of Chemical Sciences, Geosciences, and Biosciences, Office of Basic Energy Sciences of the U.S. Department of Energy.

## MATERIALS AND METHODS

### Chlamydomonas strains and media

Strains CC1009 (mt-) and CC2343 (mt+) were purchased from the *Chlamydomonas* Resource Center at the University of Minnesota. CC1009 is derived from a wild-type strain isolated G.M. Smith near Amherst, MA in 1945. CC1009 has been separated from the common c137 (CC124 and CC125) and Sager (CC1690) reference lines since the 1950s (Pröschold et al., 2005) and can assimilate nitrate. CC2343 was isolated from soil near Melbourne, FL in 1988 by Spanier (Spanier et al., 1992). Strains were maintained on agar plates containing either Tris-Acetate-Phosphate media (TAP) (Gorman and Levine, 1965), Sueoka’s High Salt media (HS) (Sueoka, 1960), or 2NBH media (Davey et al., 2012), which is a modified Bristol media containing twice the sodium nitrate. All media used Hutner’s trace elements.

### Growth, conditions, and sample collection

Strains were precultured photoautotrophically in 125 mL Erlenmeyer flasks containing 50 mL 2NBH media under constant 100 µmol photons m^-2^ s^-1^ cool white light (4000K color temperature) with 16-hour day length at room temperature (∼22°C) and bubbled constantly with sterilized ambient air (0.04% CO_2_). To maintain highly controlled conditions, cultures were grown in environmental photobioreactors (ePBR) (Lucker et al., 2014). ePBR culture column height was set to 15 cm using 330 mL 2NBH media. Preculture samples were inoculated into ePBRs to a cell density approximately 1-2 x 10^5^ cells mL^-1^, then grown for several days using standard illumination for a 14:10 h (light:dark) sinusoidal diurnal cycle with a peak light intensity of 2000 µmol photons m^-2^ s^-1^ until the culture reached a target chlorophyll concentration of 3-4 mg mL^-1^ total chlorophyll. Technical issues and time restrictions resulted in performing an additional experiment with CC2343 (CC2343 replicate #6). CC2343 replicate #6 was inoculated to a denser concentration (1-2 x 10^6^ cells mL^-1^) and grown at 1000 µmol photons m^-2^ s^-1^ constant light for 24 hours before the culture reached 3-4 mg mL^-1^ total chlorophyll (Supp. Fig. X). Culture density was maintained at 3-4 mg mL^-1^ total chlorophyll by turbidostat-controlled automatic dilution of fresh 2NBH media (Davey et al., 2012). For normoxia conditions, ePBRs were sparged with high CO_2_ gas (5% CO_2_, 21% O_2_, balanced N_2_) through a 5 mm gas dispersion stone (pore size 10-20 mm) at a flow rate of 0.35 L min^-1^ for 60 seconds every 60 minutes after being sterilized by a Whatman HEPA filter (cat. no. 6705-3602). Once cultures stabilized to their set target chlorophyll concentration, the light intensity was set to 2000 µmol photons m^-2^ s^-1^ (i.e., full sunlight) constant 24-hour light for at least 72 hours to minimize the effects of the diurnal light cycle before starting hyperoxia treatment. While 2000 µmol photons m^-2^ s^-1^ light is generally considered to be a very high light level, we would like to note in the ePBRs, cells in the lower portion of the water column are still light limited.

To induce hyperoxic conditions, the gas to the ePBRs was switched to pure oxygen enriched with CO_2_ (5% CO_2_, 95% O_2_) and sparged for 2 minutes to ensure all reactors were induced at the same time. Thereafter, gas resumed sparging 60 seconds every 60 minutes. The light was maintained at constant 24-hour 2000 µmol photons m^-2^ s^-1^. To collect samples for RNA sequencing, approximately 30 mL cell culture was withdrawn from each ePBR, transferred to a 50 mL conical tube, centrifuged at 3,000 *rcf* for 5 minutes at 4°C, flash frozen in liquid nitrogen, and stored at -70°C until ready for extraction. Samples for RNA-seq were collected at 0’ (before gas swap), 1, 3, 6, 12, 24, and 48 hours after hyperoxia induction.

### Chlorophyll measurements

Chlorophyll content was measured using methods developed by Porra (Porra et al., 1989; Porra, 2002). The extraction buffer was modified from 80% acetone to a solution of 60% acetone and 40% dimethyl sulfoxide (DMSO) as performed by Neofotis (Neofotis et al., 2021).

### Comparative genomics

Genomic DNA was isolated as described by Fawley and Fawley (Fawley and Fawley, 2004) and purified using an MPBio GENECLEAN Turbo Kit (Cat. No. 111102-600) following the manufacturer’s instructions. DNA sequencing was performed by the Michigan State University Research Technology Support Facility (RTSF) using the Illumina HiSeq 2500 platform to generate 125 bp paired end reads. Raw reads were quality-checked using FastQC version 0.12.1 using default settings (bioinformatics.babraham.ac.uk/projects/fastqc/). Adapters and low-quality bases were trimmed with Trimmomatic version v0.39 using settings LEADING:10 TRAILING:10 SLIDINGWINDOW:4:20 MINLEN:75 (Bolger et al., 2014). Trimmed reads were aligned using bwa-mem2 version 2.2.1 with default settings (Md et al., 2019). Aligned reads were sorted and indexed using samtools version 1.19.2 with default settings (Li et al., 2009). Variants were called and a consensus sequence generated using bcftools version 1.22. When the *call* command was issued the --ploidy 1 flag was used to indicate a haploid genome. Ecotype consensus sequences were aligned to the *Chlamydomonas* reference genome v6.0 and to each other using the *nucmer* alignment tool (flags --maxmatch -c 100 -b 500 -l 50) in the MUMmer4 package version 4.0 (Marçais et al., 2018). Genomes were compared using the *dnadiff* tool.

### RNA isolation and sequencing

Total RNA was extracted using a Qiagen RNeasy Plant Mini Kit (cat. no. 74904) by manufacturer’s instructions. Total RNA quantification was performed on a Qubit 1.0 Fluorometer using Qubit RNA High Sensitivity Assay Kit (cat. no. Q32852). RNA sample integrity was verified using an RNA 6000 Pico Assay chip and an Agilent 2100 Bioanalyzer. The Michigan State University RTSF prepared the RNA-sequencing libraries using the Illumina TruSeq RNA Library Prep Kit generating approximately 42 million uniquely mapped 125 bp paired end reads per sample.

### Transcriptome data analysis

RNA-seq reads were aligned to the *Chlamydomonas* reference genome v6.1 (Craig et al., 2023) with STAR 2.7.10a using default parameters (Dobin et al., 2013). Alignment results were sorted using SAMtools 1.15 (Li et al., 2009) and reads were counted using HTSeq (Anders et al., 2015). HTSeq read count files were analyzed using R and the DESeq2 package (Love et al., 2014). Genes with fewer than 10 read counts in at least 5 samples were removed. Principal component analysis (PCA) was performed using the DESeq2 “plotPCA” function. During differential gene expression analysis, the DESeq2 “lfcshrink” function was used with the “*apeglm”* shrinkage estimator to minimize the number false positive hits (Zhu et al., 2018).

Genes with a log_2_ fold-change (LFC) > 1 and an adjusted p-value < 0.05 were defined as significant. Additional analysis and visualization were performed using Python, Jupyter Notebooks, and the SciPy package (Virtanen et al., 2020). Pearson correlation analysis was performed using SciPy library “pearsonr” function. Expression values were normalized to fragments per kilobase of transcript per million reads mapped (FPKM) using gene mRNA lengths (Zhao et al., 2021).

### Differential gene expression analysis

We utilized an established RNA-seq analysis pipeline using DESeq2 to identify significant differentially expressed genes (DEG). Genes with fewer than 10 counts in at least 3 samples were excluded. To reduce the number of false positive hits due to high expression variance or a low number of counts, we employed the *apeglm* shrinkage estimator using the “lfcshrink” DESeq2 function (Zhu et al., 2018). Genes with a LFC > 1 and an adjusted p-value < 0.05 when compared to the 0’ time point were classified as significant DEGs.

### Functional enrichment analysis

Differentially expressed genes displaying a LFC > 1 and an adjusted p-value < 0.05 for each ecotype and timepoint were selected for functional enrichment analysis. Each gene set was input into the Algal Functional Annotation Tool (Lopez et al., 2011) to identify gene ontology (GO) term, MapMan ontology, and KEGG pathway enrichment. Terms with less than two hits and a score greater than 0.05 were filtered out. The results were exported for visualization in a Jupyter Notebook using Seaborn (Waskom, 2021).

### Literature comparison

Sequence data were retrieved from the NCBI SRA database, specifically using SRP009466 (Fang et al., 2012) and PRJNA1123657 (Pancheri et al., 2024). Reads were aligned to the *Chlamydomonas reinhardtti* genome v6.1 (Craig et al., 2023) and filtered as described in the “RNA isolation and sequencing” methods subsection. DEGs were identified as described in the “Differential gene expression analysis” methods subsection using the same LFC and adjusted p-value thresholds. The set of low CO_2_ genes are defined as DEG from the wild-type low (L) or very low (VL) samples. The set of singlet oxygen genes are defined as genes that appear in both the wild-type liquid rose Bengal (RB) and liquid high light (LWTHL) samples.

