## Supplemental Figures and Legends for "Natural genetic variation reveals divergent transcriptomic responses to hyperoxia in two *Chlamydomonas reinhardtii* ecotypes"

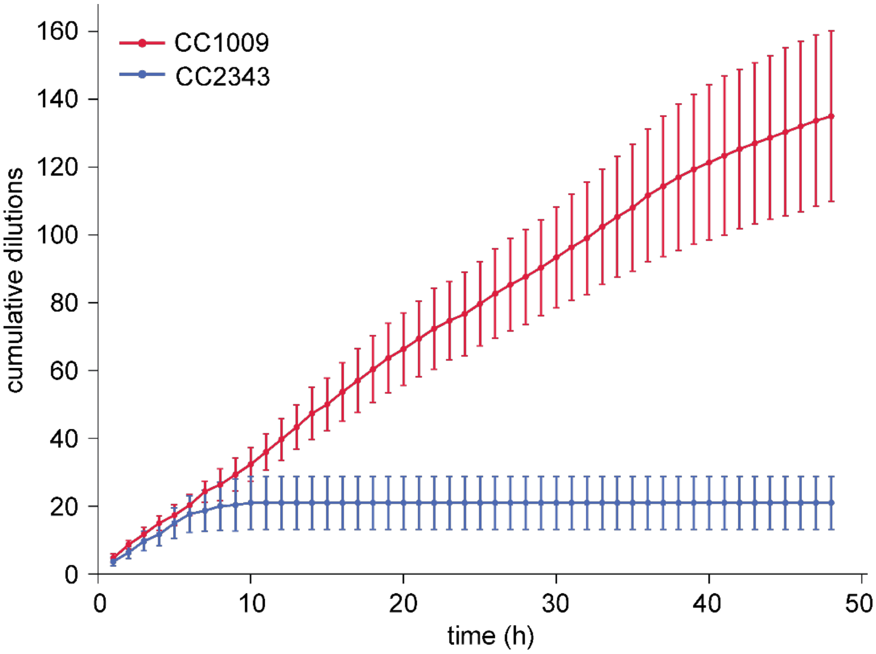


**Figure S1. Cumulative ePBR turbidostat dilutions under hyperoxia.** Data represents mean ± SD of 3 biological replicates strains CC1009 (red) and CC2343 (blue).

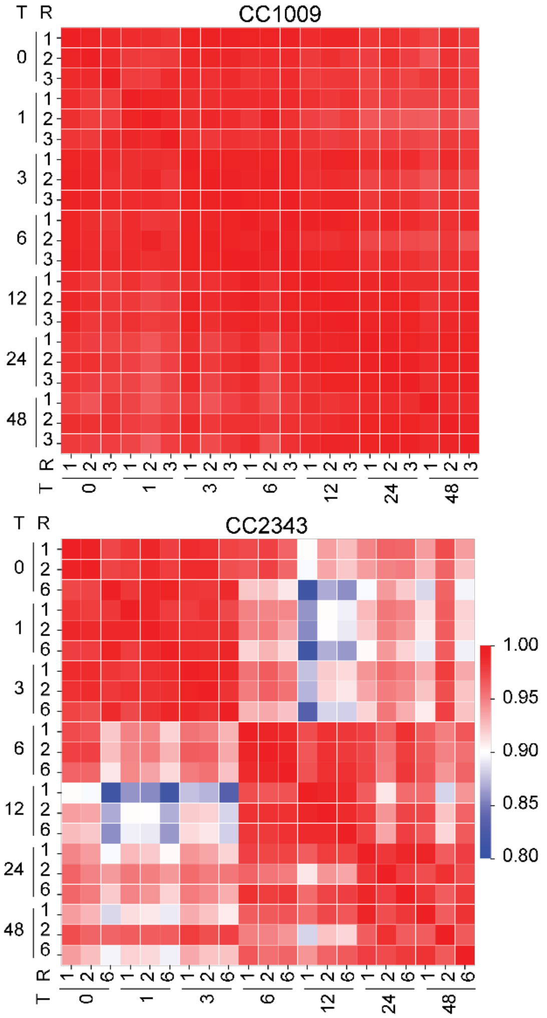


**Figure S2. Pearson correlation analysis across timepoints.** Heatmap shows Pearson correlation coefficients between all samples for ecotype CC1009 (left) and CC2343 (right). The color scale represents correlation values ranging from 0.8 (blue) to 1.0 (red). T indicates time point after hyperoxia treatment (h), R indicates replicate number.

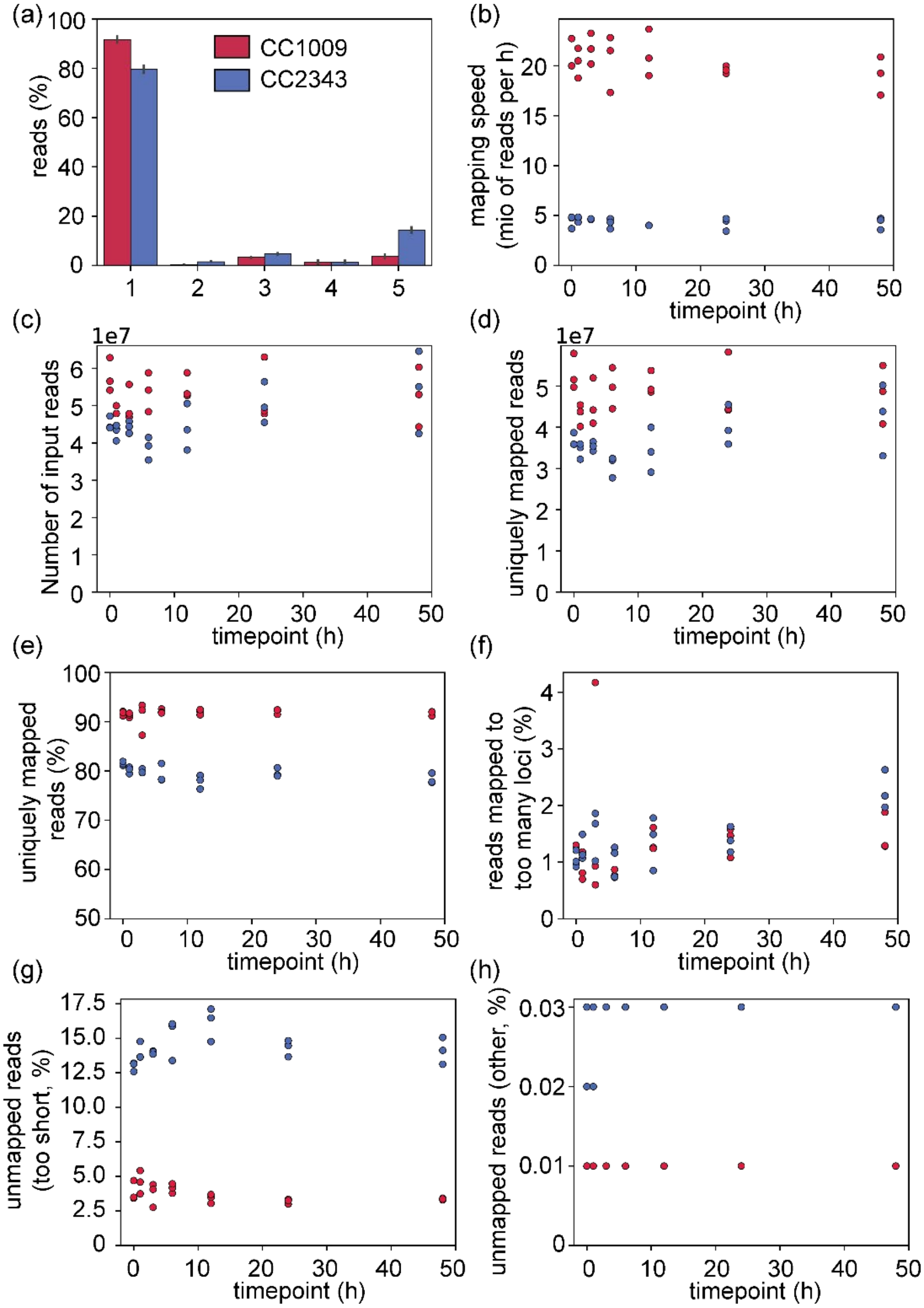


**Figure S3. RNA-seq processing metadata analysis.** (a) Mean ± SD percent reads across different mapping categories for strains CC1009 (red) and CC2343 (blue) of 3 biological replicates (1 = unique mapped reads, 2 = mismatch rate, 3 = multiple loci, 4 = too many loci, 5 = too short), (b) mapping speed (millions of reads per hour), (c) number of input reads, (d) number of uniquely mapped reads, (e) percentage of uniquely mapped reads, (f) percentage of reads mapped to too many loci, (g) percentage of reads unmapped due to being "too short", (h) percentage of reads unmapped for other reasons. Data are shown as individual replicates in subplots b-h for strains CC1009 (red) and CC2343 (blue).

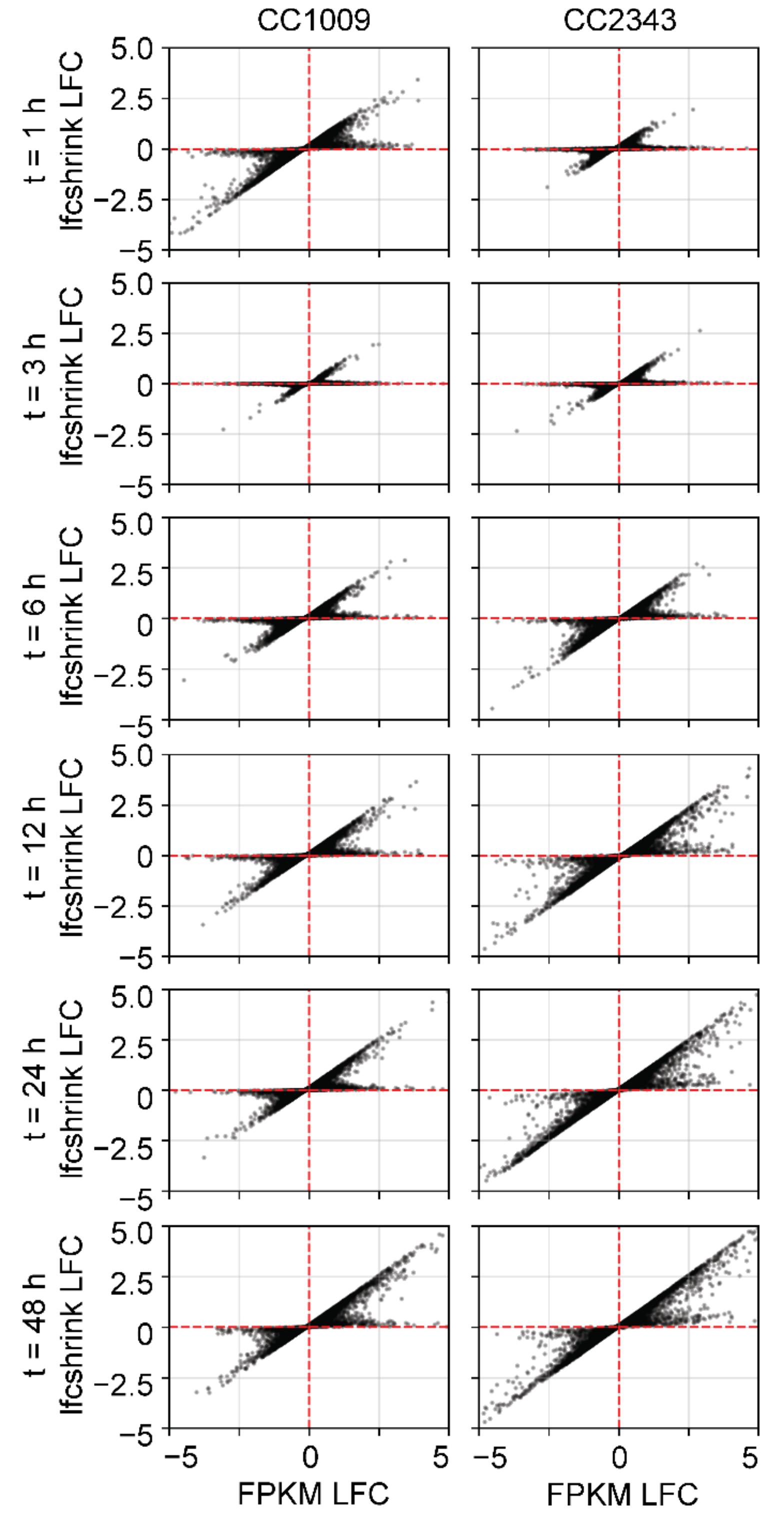


**Figure S4. Comparison of gene expression fold changes before and after *lfcshrink* application.** The figure displays the $\log_2$ fold change (LFC) of all genes (grey/black dots) for strains CC1009 (left column) and CC2343 (right column) across different timepoints as indicated (rows). Each plot compares the pre-shrinkage LFC (x-axis) and post-shrinkage LFC (y-axis). The dashed red lines indicate a fold change of 0.

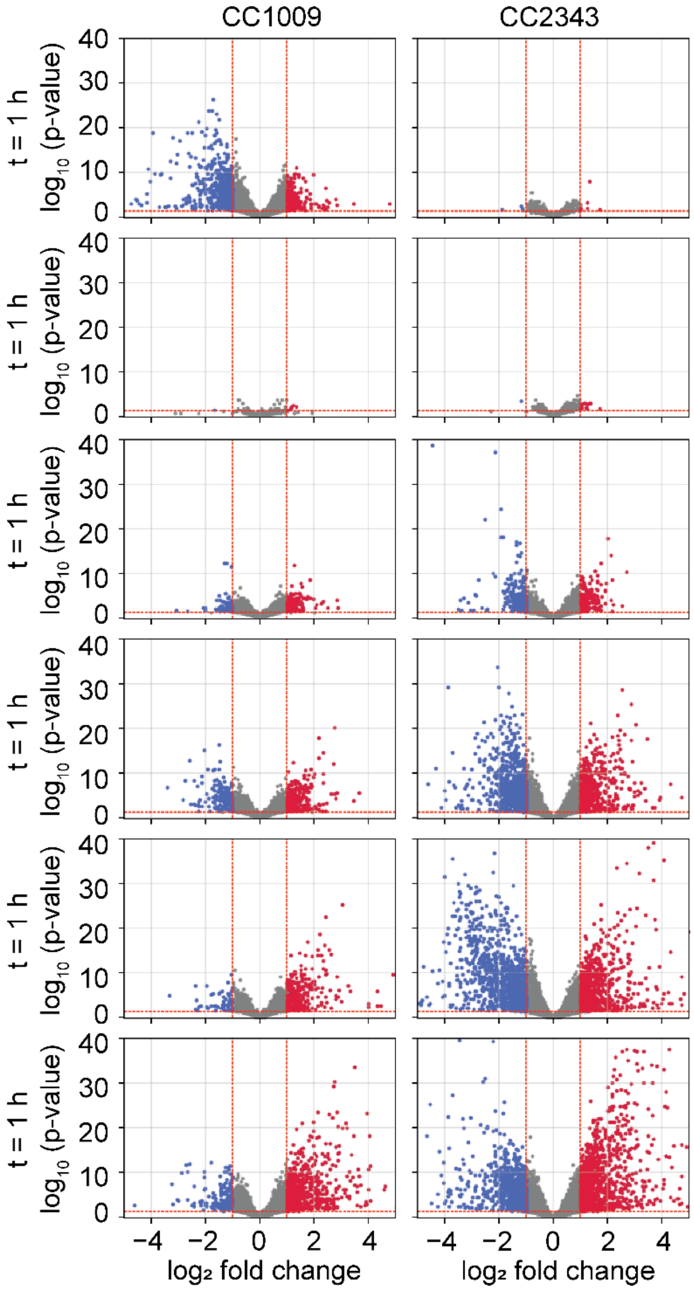


**Figure S5. Volcano plots show differential gene expression over time.** The figure displays the $log_2$ fold change (LFC) versus statistical significance ($-log_{10}$ adjusted p-value) for strains CC1009 (left column) and CC2343 (right column). Each row represents a different timepoint. Genes are colored by significance: red (upregulated), blue (downregulated), and grey (non-significant). DEGs are defined genes with a LFC > 1 and an adjusted p-value < 0.05.

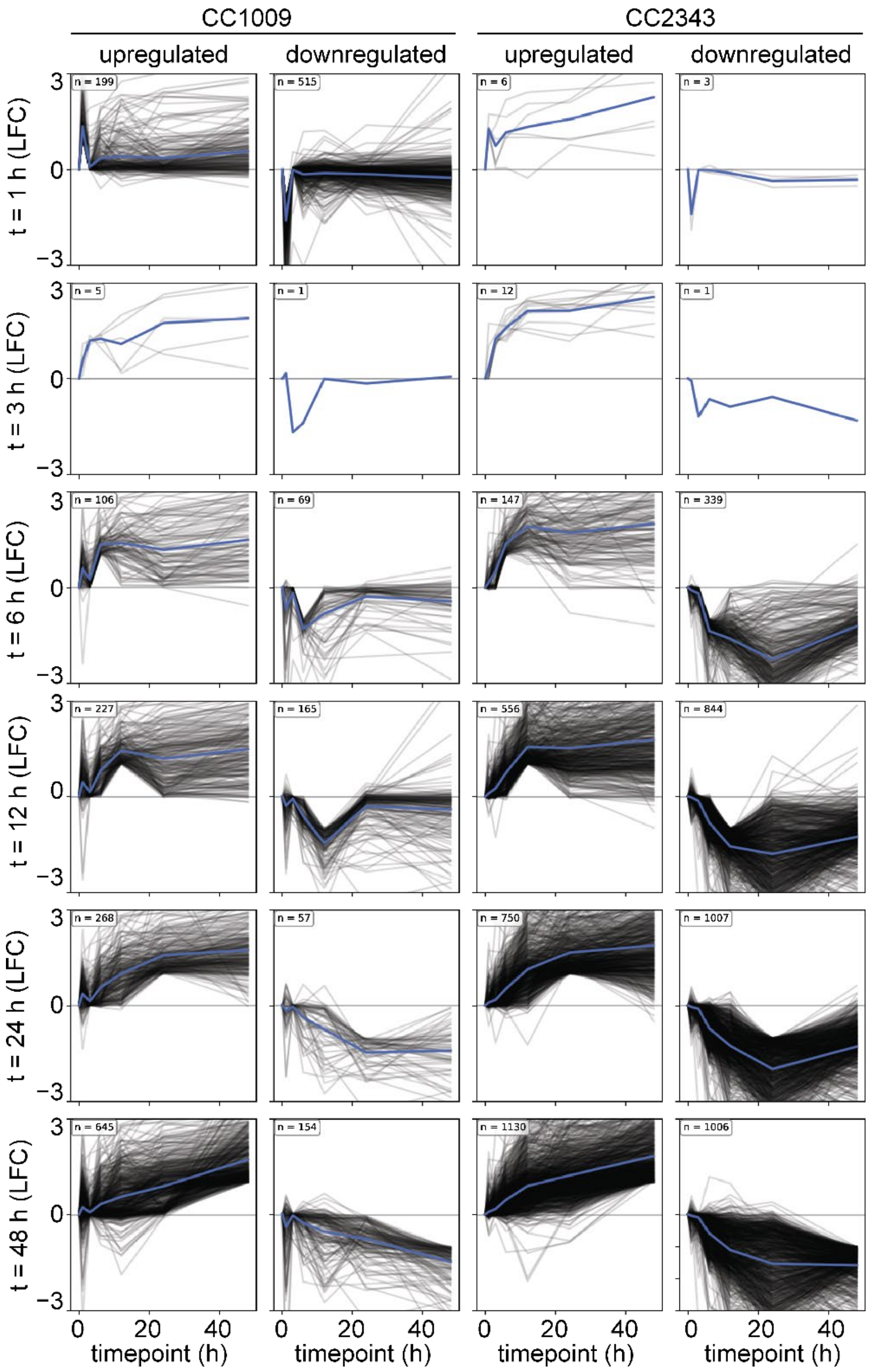


**Figure S6. Gene expression profiles for differentially expressed genes (DEGs) over time**. The figure displays the $\log_2$ fold change of upregulated and downregulated genes for strains CC1009 (left two columns) and CC2343 (right two columns). Each row represents a different timepoint. Individual genes are shown in black/grey, with mean expression across all genes highlighted in blue. DEGs were defined as genes with a LFC > 1 and an adjusted p-value < 0.05.

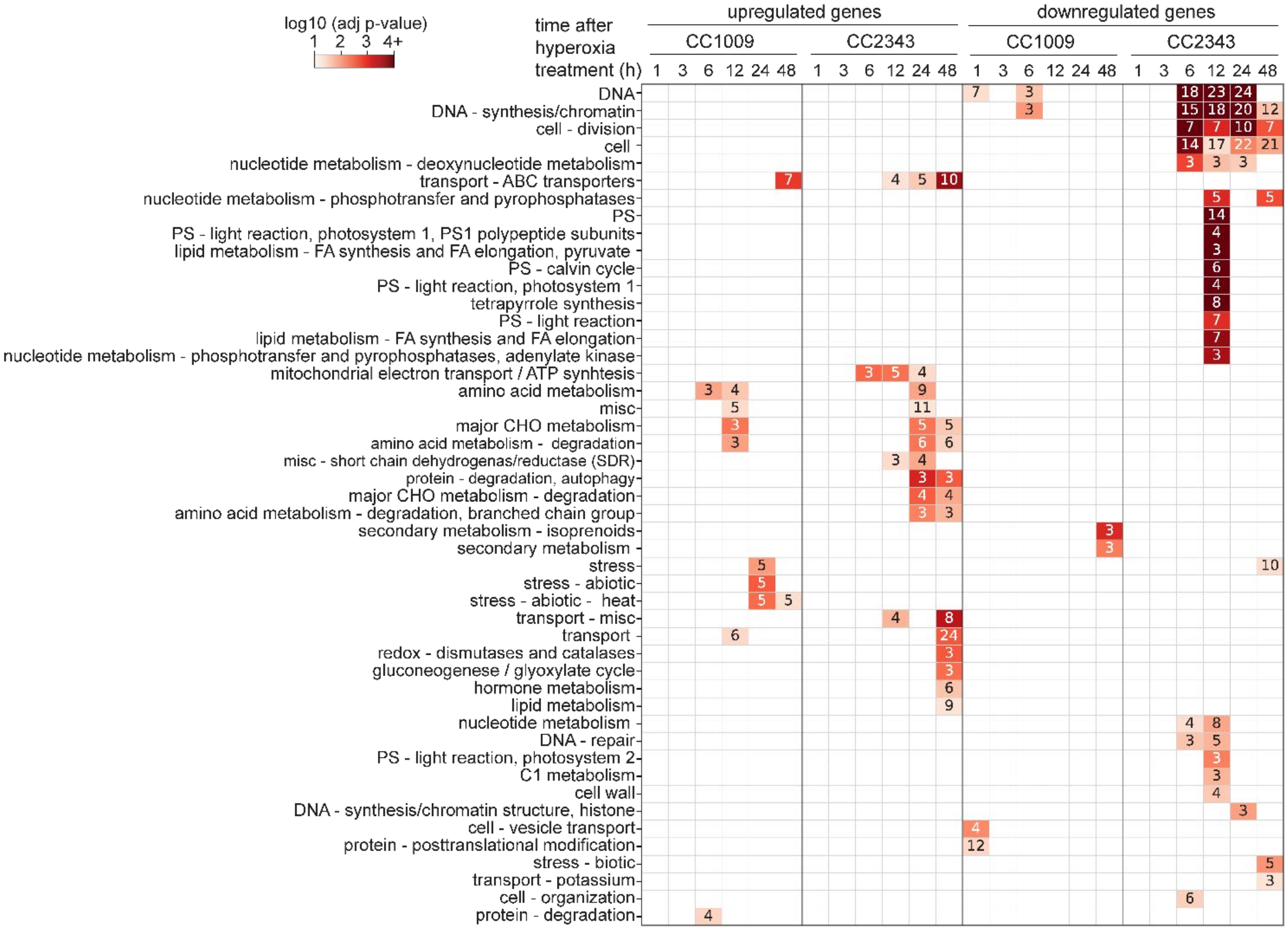


**Figure S7. Functional enrichment analysis of MapMan terms.** Heatmap showing the number of significantly enriched functional categories for strains CC1009 and CC2343 across different timepoints. The numerical values in each cell represent the number of genes (hits) associated with that term. The color scale represents the $-\log_{10}$(p-value), ranging from 1 (light red) to > 4 (dark red). Only terms > 2 hits and a p-value < 0.05 are displayed.

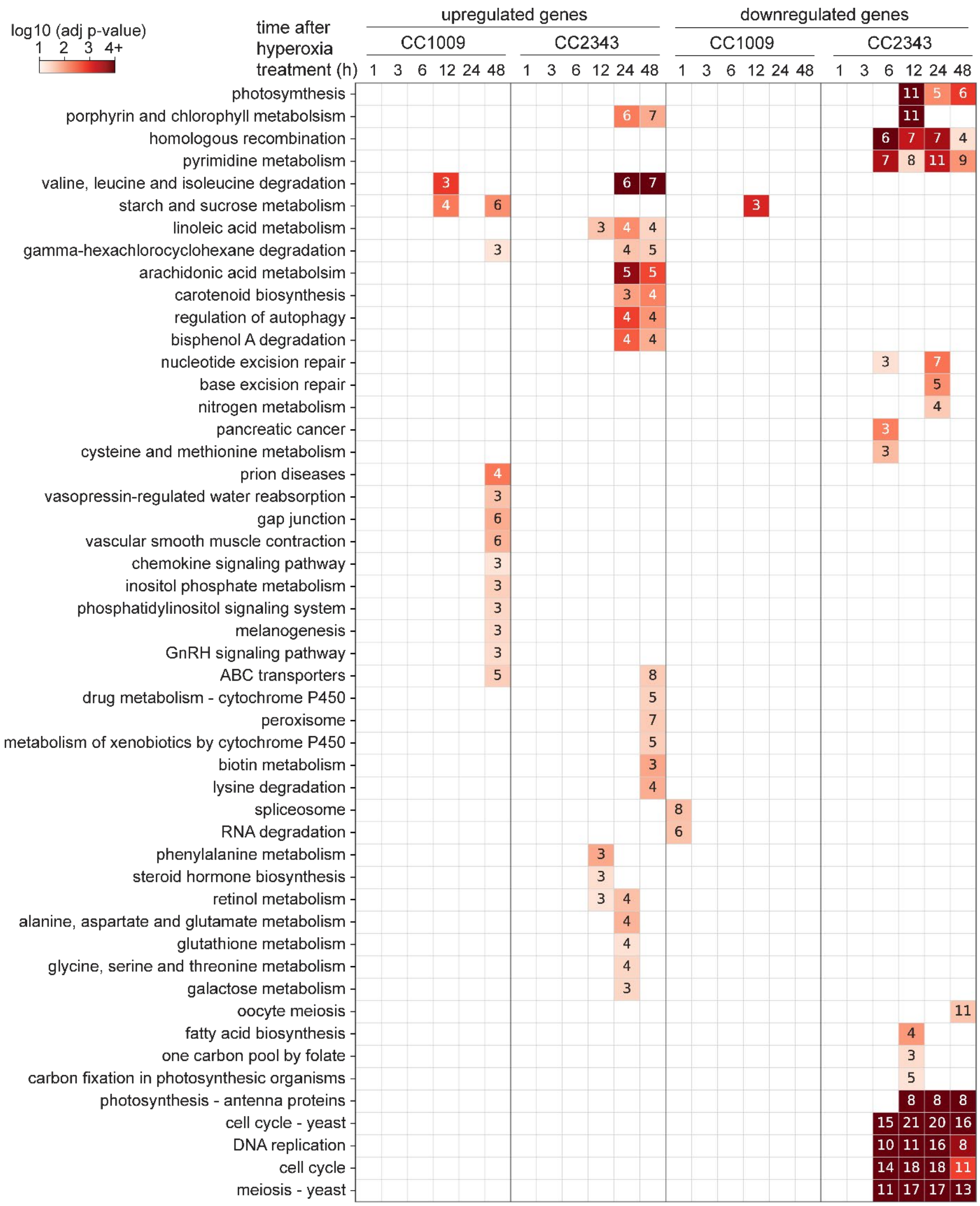


**Figure S8. Functional enrichment analysis of KEGG pathways.** Heatmap showing the number of significantly enriched functional categories for strains CC1009 and CC2343 across different timepoints. The numerical values in each cell represent the number of genes (hits) associated with that term. The color scale represents the $-\log_{10}$(p-value), ranging from 1 (light red) to > 4 (dark red). Only terms > 2 hits and a p-value < 0.05 are displayed.

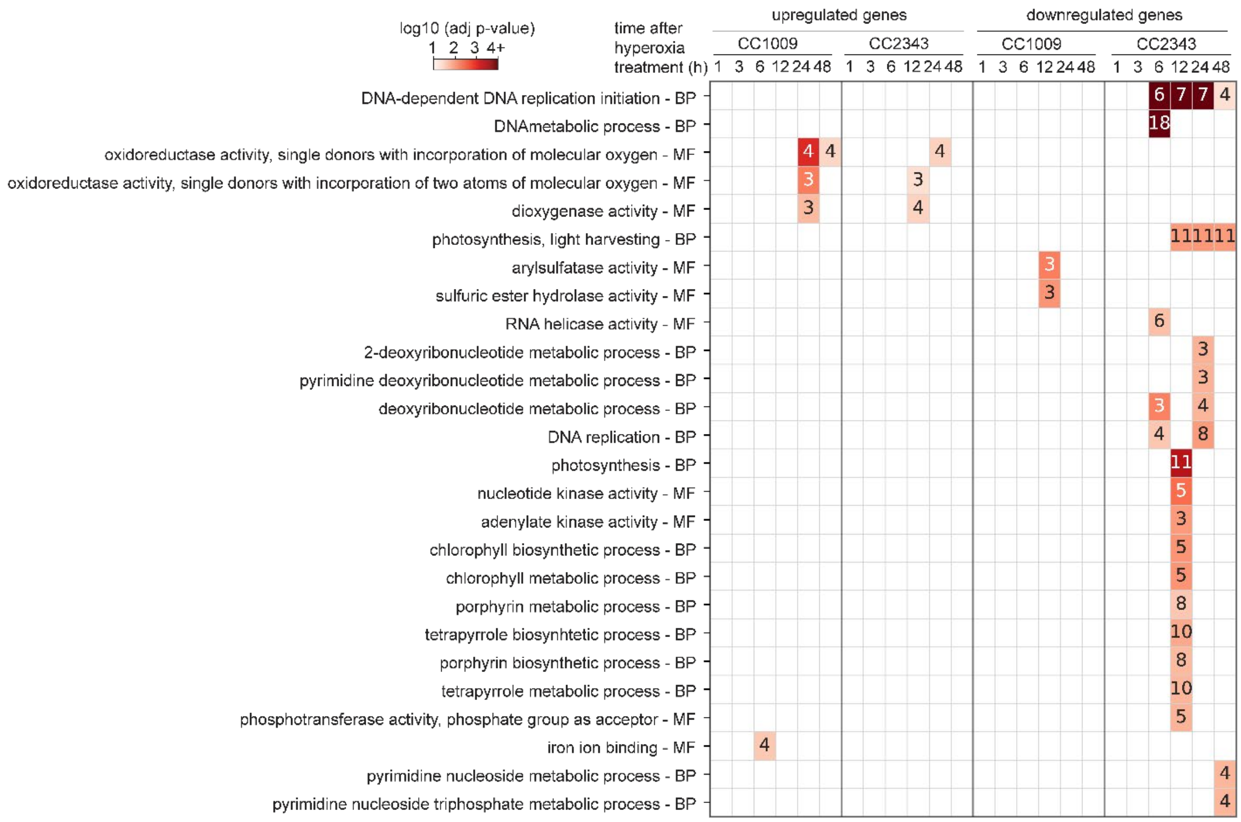


**Figure S9. Functional enrichment analysis of Gene Ontology (GO) terms.** Heatmap showing the number of significantly enriched functional categories for strains CC1009 and CC2343 across different timepoints. The numerical values in each cell represent the number of genes (hits) associated with that term. The GO terms are categorized into BP (Biological Process) or MF (Molecular Function), as indicated in the row labels. The color scale represents the -$\log_{10}$(p-value), ranging from 1 (light red) to > 4 (dark red). Only terms > 2 hits and a p-value < 0.05 are displayed.
