## Supplementary material for "Natural genetic variation reveals divergent transcriptomic responses to hyperoxia in two *Chlamydomonas reinhardtii* ecotypes": Table S1

| Query Genome | CC1009 | CC2343 | CC2343 |
|---|---|---|---|
| **Reference Genome** | **CC4532** | **CC4532** | **CC1009** |
| **Alignments** | | | |
| 1-to-1 | 211 | 158 | 154 |
| AvgIdentity | 99.5199 | 95.9329 | 95.8171 |
| M-to-M | 227 | 227 | 227 |
| AvgIdentity | 99.5191 | 95.9434 | 95.8296 |
| **Feature Estimates** | | | |
| Breakpoints | 333 | 332 | 332 |
| Relocations | 5 | 7 | 7 |
| Translocations | 0 | 0 | 0 |
| Inversions | 0 | 0 | 0 |
| Insertions | 121 | 100 | 98 |
| TandemIns | 74 | 17 | 12 |
| **SNPs** | | | |
| TotalSNPs | 370576 | 3295794 | 3409471 |
| TotalGSNPs | 74622 | 519400 | 533290 |
| TotalIndels | 178020 | 1200180 | 1304192 |
| TotalGIndels | 3285 | 14900 | 15739 |

**Table S1. Genomic comparison summary using dnadiff.** The table summarizes alignment statistics, feature estimates (e.g., breakpoints and insertions), and SNP/Indel counts. Each column represents a unique query-reference pair as reported in the dnadiff output files.
